# Maintaining grass coverage increases methane uptake in Amazonian pasture soils

**DOI:** 10.1101/2021.04.26.441496

**Authors:** Leandro Fonseca de Souza, Dasiel Obregon Alvarez, Luiz A. Domeignoz-Horta, Fabio Vitorino Gomes, Cassio de Souza Almeida, Luis Fernando Merloti, Lucas William Mendes, Fernando Dini Andreote, Brendan J. M. Bohannan, Jorge M. Rodrigues, Klaus Nüsslein, Siu Mui Tsai

## Abstract

Cattle ranching is the largest driver of deforestation in the Brazilian Amazon. The rainforest- to-pasture conversion affects the methane cycle in upland soils, changing it from sink to source of atmospheric methane. However, it remains unknown if management practices could reduce the impact of land-use on methane cycling. In this work, we evaluated how pasture management can regulate the soil methane cycle either by maintaining continuous grass coverage on pasture soils, or by liming the soil to amend acidity. Methane fluxes from forest and pasture soils were evaluated in moisture-controlled greenhouse experiments with and without grass cover (*Urochloa brizantha* cv. Marandu) or liming. In parallel, we assessed changes in the soil microbial community structure of both bare pasture soil as well as rhizosphere soil through high throughput sequencing of the 16S rRNA gene, and quantified the methane cycling microbiota by their respective marker genes related to methane generation (*mcr*A) or oxidation (*pmo*A). The experiments used soils from eastern and western Amazonia, and concurrent field studies allowed us to confirm greenhouse data. The presence of a grass cover not only increased methane uptake by up to 35% in pasture soils, but also reduced the abundance of the methane-producing community. In the grass rhizosphere this reduction was up to 10-fold. Methane-producing archaea belonged to the genera *Methanosarcina sp*., *Methanocella sp*., *Methanobacterium sp*., and Rice Cluster I. Further, we showed that liming compromised the capacity of forest and pasture soils to be a sink for methane, and instead converted formerly methane-consuming forest soils to become methane sources in only 40-80 days. Our results demonstrate that pasture management that maintains grass coverage can mitigate soil methane emissions, if compared to a bare pasture soil.

## Introduction

The establishment of pasture lands is the main cause of deforestation in the Amazon region (Dias et al., 2016; Margulis, 2003). This transformation of rainforest into pastures leads to the net increased emission of the powerful greenhouse gas methane, and turns a methane consuming forest soil into a methane producing pasture soil (Meyer et al., 2020; Fernandes et al., 2002; Steudler et al., 1996; Goreau; De Mello, 1988). The resulting greenhouse gas emissions account for half of Brazil’s greenhouse gas production, and have already exceeded national emissions from fossil fuel by more than 20% (Bustamante et al., 2012; Fearnside & Imbrozio Barbosa, 1998). Livestock production is responsible for the emission of several greenhouse gases, and current studies work on minimizing this impact (Herrero et al. 2016, Fiqueiredo et al. 2017). Recently, methane emissions from soil have become a focus of investigations because they might be agriculturally manageable. The impact of land-use conversion on the annual balance of gas fluxes is noticeable considering pastures in western Amazonia can emit up to 270 mg C-CH_4_/ m^2^, while nearby forest soils can consume up to 470 mg C-CH_4_/ m^2^ (Steudler et al., 1996).

Methane (CH_4_) gas has an 86-fold greater potential to retain heat in the atmosphere compared to that of CO_2_, calculated over a 20-year period (IPCC, 2013). The global methane emissions are mainly driven by human activities such as livestock, irrigated agriculture, oil and gas production, and landfill decomposition (IPCC, 2013). Soil methane cycling is strongly dependent on the microbiota, since the biogenic source of this gas are methanogenic archaea. The biological consumption of methane is controlled by methanotrophs, mostly bacteria. In soil, the balance between methanotrophic bacteria and methanogenic archaea is related to environmental conditions (i.e., moisture, temperature, soil density, and pH) and is sensitive to changes in agricultural management (Le Mer & Roger, 2001; Liu et al., 2007; Tian et al., 2015).

Methanotrophic bacteria in the soil are Gram-negative, belonging to *Gammaproteobacteria* and *Alphaproteobacteria, Verrucomicrobia*, and candidates in the phylum NC10 (Hanson & Hanson, 1996; Knief, 2015, Ettwig et al. 2010). The initial step of methane oxidation occurs through its conversion to methanol, which is mediated by the enzyme methane monooxygenase (MMO). Methanogenic archaea traditionally comprise members from eight orders within the phylum *Euryarcheota*: *Methanopyrales, Methanobacteriales, Methanococcales, Methanomicrobiales, Methanocellales, Methanosarcinales, Methanomassiliicoccales*, and ‘*Candidatus* Methanophagales’ (Evans et al. 2019), with additional candidates in the phylum *Bathyarchaeota* (Kallistova et al., 2017). Methanogenesis is controlled by archaea and is the final step in an anaerobic pathway that begins with the hydrolysis of organic polymers, fermentation of the resulting monomers and of initial fermentation products, and ends up in the production of CH_4_ mostly from acetate, hydrogen, and CO_2_. The final step in the methanogenesis pathway is facilitated through the action of the enzyme methyl-coenzyme M reductase, coded by the *mcrA* gene, which can be used as a methanogen-specific marker for molecular studies (Serrano-Silva et al., 2014). The ability to produce methane has recently been demonstrated in cyanobacteria and plants, however, it is believed that this is a by-product from reactions of photosynthesis (Bižić et al., 2020; Keppler et al., 2006).

The relationship between changes in land-use and the response of the soil microbial community is not well understood (Nazaries et al., 2013; Tate, 2015), but previous studies have shown significant impacts on microbial diversity in the Amazon region (de Carvalho et al., 2016; Mendes et al., 2015; Navarrete et al., 2015; Rodrigues et al., 2013; Jesus et al., 2009). In well-managed pastures in the Amazon region, the grass root system can redistribute carbon to deeper layers, where it is less susceptible to decomposition (Fearnside & Imbrozio Barbosa, 1998). On the other hand, we can expect that degraded pastures with large bare soil areas can facilitate the release of carbon from the system, with superficial grassroots, higher loss of soil organic matter, and lower carbon stocks (Segnini et al. 2019). Proper management of pasture may involve several practices, such as soil acidity correction and continuous maintenance of grass cover to protect soil from erosion. These practices are particularly important in the Amazon region given the environmental extremes of this area. Altogether, the high soil acidity, high rainfall, and high temperatures combined with exposure of the soil to equatorial solar radiation constitute factors that are associated with increasing erosion and soil degradation (Demattê and Demattê, 1993). Soil degradation is the long-term decline in the soil’s productivity and its environment moderating capacity, with soil quality loss and reduction in attributes related to specific functions of value to humans (Lal, 2001).

The effect of soil liming on methane fluxes is still poorly understood, and studies of temperate forests show that liming can lead to both an increase and a decrease of methane consumption (Wang et al. 2021; Borken and Brumme, 1997; Butterbach-Bahl et al., 2002). In wheat-focused agriculture liming has led to an increased consumption of methane in soil (Hütsch et al., 1994). The increased methane consumption after liming was also observed in Mediterranean semiarid soils under lupine, wheat, and triticale, a hybrid of wheat and rye (Barton et al., 2013; García-Marco et al., 2016). However, for tropical soils little information is available regarding what influence liming has on methane production and consumption. In an assessment of greenhouse gas fluxes from soils under soybean cultivation in Brazil, the acidity correction presented no effect on methane fluxes (Lammel et al., 2018). Likewise, in a field experiment in Puerto Rico, soil consumption of atmospheric CH_4_ in an intentionally acidified soil was about one-fourth of that at pH 6, and was not restored after liming (Mosier et al., 1998).

There is a growing consensus that the key to understand major soil functions lies where plants and soil meet, in the rhizosphere (Lau et al., 2011). The rhizosphere is a micro-environment with differentiated soil conditions and steep overlapping gradients, in which pH can be up to 2 units more acidic or more basic than the soil surrounding the rhizosphere. The rhizosphere can present heterogeneous concentrations of oxygen and moisture and can be enriched in root exudates like sugars and organic acids (Philippot et al., 2013). These factors affect soil methane cycling not only by providing organic substrate for methanogenesis but also by promoting the oxidation of methane in the rhizosphere. Despite its known role in flooded rice soils (Frenzel et al., 1992), little is known about the impact of the rhizosphere on methane cycling in upland soils since these soils are commonly considered to be a methane sink, not a source (Philippot et al., 2009). Thus, improved understanding of how the rhizosphere of land-intensive tropical pastures affects soil methane cycling can yield new strategies to mitigate greenhouse gas emissions related to cattle ranching.

The intensive land use in agriculture and cattle ranching in Amazonia leads to soil and pasture degradation ranging from 50% to 70% of total area (Dias-Filho, 2017). Within this context this research aims to evaluate how the management of pastures can affect soil methane cycling. We hypothesized that CH_4_ production is reduced by liming soils and by continuous grass coverage due to the influence of the rhizosphere of *Urochloa brizantha* cv. Marandu, a grass widely used in pastures in Brazil. To test this hypothesis, we measured methane fluxes in soils from pasture field sites with and without grass cover, and compared the microbiota in bare pasture soil to that of the rhizosphere of grass covered soil. These studies were complemented with greenhouse experiments where soil acidity was adjusted and grass was planted, gas flux rates of the soil-air CH_4_-fluxes were measured, and shifts in the soil microbial community between bare soil and the rhizosphere of *Urochloa brizantha* cv. Marandu were determined.

## Materials and Methods

### Sampling

These studies were performed with soils from both western and eastern Amazonia. In the western region (hereafter “Ariquemes”) sampling was carried out in April of 2017 at the Fazenda Nova Vida near Ariquemes, RO (10°10’49.5” S, 62°49’23.9” W). While in the eastern region (hereafter “Tapajós”), the samples were taken at the National Forest of Tapajós and the immediately surrounding areas near Belterra, PA (3°07’53.8” S, 54°57’24.2” W), in August of 2019. The sampled soils were used in two rounds of greenhouse experiments at the Center for Nuclear Energy in Agriculture, SP, Brazil (22°42’27.7” S, 47°38’41.0” W). In addition to the soil sampling a field study was performed, but only in the Tapajós region (detailed below).

Western Amazonia has been studied for the impacts of conversion from forest to pasture, with extensive scientific literature characterizing ecosystem responses to conversion (de Moraes et al., 1996; Herpin et al., 2002; Reiners et al., 1994) for representing a region with a high degree of exploitation. The Fazenda Nova Vida region has fragments of primary forest and pastures of different ages. The sampled pasture area was established in 1972, and since then managed by cattle rotation, with the use of fire only to control eventual pests, mechanical removal of invasive trees, and at least one record of liming 15 years before the sampling. Soils sampled varied from average clay to sandy texture.

Eastern Amazonia represents areas of more recent exploration. The Tapajós National Forest was sampled as a model of a conservation area and the pasture chosen is in a small property in Belterra, PA. The pasture used here was established between 1989-1994 and supported cattle at the time of sampling, had sparse signs of degradation, fire was applied when necessary to control invasive plants, and it has no history of liming. Soils sampled varied from average clay to sandy texture.

During each expedition, we sampled 20-30 kg of soil from the upper 0-10 cm layer of 5 equidistant sampling points along a linear gradient of 200 m at each site, from areas under Primary Forest and Pastures with *Urochloa brizantha* cv. Marandu. The sampled soils were transported to the Cell and Molecular Biology Laboratory at the University of Sao Paulo, CENA-USP, with fresh samples for chemical analysis and greenhouse experiments, or were frozen at the end of each sampling day and stored at −20 °C for future molecular analyses.

### Field Study

At two pastures in the Tapajós region, 100 m side squares were established and 4 points in the square corners, plus a point in the center, were selected to evaluate CH _4_ fluxes and to sample soils for molecular and chemical analysis. Those 5 points had grass coverage at the time, and prior to gas flux measurements with static gas collection chambers the grass leaves were cut to their stems (2 cm above soil surface) and removed. Following chamber removal, the roots were collected, and the rhizospheric soil was sampled and stored at −20 °C. Adjacent to each of the five sampling points we had selected, one square meter large areas without grass (bare soil) we used to measure methane fluxes and to collect soil samples for molecular and chemical analysis.

### Greenhouse experiments

Sampled soil was homogenized, sieved (5 mm), and placed in clay pots with a capacity of 10 liters, resulting in 10 cm high soil columns with 5 kg of soil per pot. The grass was raised from seed in a subsample of the soil, and mature plants were transferred to the experimental clay pots at least 40 days after soil liming. The liming was performed by the addition of CaCO _3_ to reach pH 6.5 (water), calculated for a base saturation of 70-75%. For each treatment, four pots were used to grow *Urochloa brizantha* cv. Marandu with 4 additional pots as no-plant controls (bare soil), both at natural pH and with limed soils (4 pots x 2 soil types x 2 pH situations x 2 plant situations). At the beginning of the experiment soil moisture was standardized to ∼70% of the water retention capacity of the soil and adjusted every two to four days, taking as reference the weight variation after drying soil samples for 48 h at 75 °C. In the experiment with soil from Ariquemes, the plants were removed when they reached approximately 35 cm in height, and by shaking and with the help of a sterilized brush the rhizosphere soil was collected. Here we defined the rhizosphere as soil that remained attached to the roots even after vigorous plant shaking.

### Determination of methane fluxes in soil

The measurements of CH_4_ fluxes in both the field sampling sites and the greenhouse experiments were carried out using static gas collection chambers (20 cm in diameter, ∼6 L inner volume). Over a period of 10 minutes measurements were taken at 10 second intervals using a portable gas analyzer (UGGA, Los Gatos Research, San Jose, CA, USA). Daily flux of gases was estimated from the concentration in the chamber headspace. Daily flux (F, mass of gas m^-2^.day^-1^) was computed as (Ussiri, Lal and Jarecki, 2009):

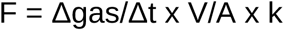

Where Δ*gas*/Δ*t* is the rate of change in CH_4_ concentration inside the chamber (i.e. mg CH_4_-C); *V* is the chamber volume (m^3^); *A* is the surface area circumscribed by the chamber (m^2^) and *k* is the time conversion factor (1440 min day^-1^). The cumulative gas emissions were calculated by linear interpolation of average emissions between two successive measurements and the sum of the results obtained over the entire study period. Finally, the data was expressed as differences in the cumulative CH_4_ fluxes in relationship to the controls. From this, we subtracted from the accumulated fluxes in the respective treatments (liming, grass coverage, and liming plus grass coverage) the control measurements of the average accumulated flux (bare soils).

### Characterization of soil chemical properties

About 600 g of soil were analyzed for their physical-chemical properties at the Laboratory of Chemical Analysis in the Soil Science Department at the “Luiz de Queiroz School of Agriculture” (ESALQ / USP) (detailed in van Raij et al., 2001). The soil attributes measured were: pH in CaCl_2_; concentrations of phosphorus, potassium, calcium, and magnesium by extraction with ion exchange resin; aluminum by extraction of potassium chloride at 1 mol/ L; potential acidity estimated by pH-SMP buffer test; organic matter by the dichromate-titrimetric method; boron by extraction with hot water; copper, iron, manganese and zinc extracted by the DTPA-TEA extractor (pH 7.3); and by calculating the sum of bases (BS); cation exchange capacity (CEC); base saturation (V%), and aluminum saturation (m%).

### DNA extraction

DNA was extracted for molecular analyses from greenhouse soils that originated in Ariquemes and Belterra (Tapajós region) and from soils of the field study in the Tapajós regions. Total DNA was extracted from soil samples using the PowerLyzer PowerSoil DNA Isolation Kit (Qiagen, Hilden, Germany) from 250 mg of soil, according to the protocol provided by the manufacturer, except that after adding solution C1 the stirring time was extended to 15 minutes followed by 3 min centrifugation (Venturini et al., 2020). The amount and quality of the DNA extracted was analyzed in a Nanodrop 2000c spectrophotometer (Thermo Fisher Scientific, Waltham, MA, USA) at an optical density of 260 nm. The total DNA extracted was stored at −20 °C.

### Abundance of methane producers and oxidizers

Real-time quantitative PCR (qPCR) was used to quantify the genes associated with methane cycling *mcrA and pmoA* (Table S1) in total soil DNA samples. For each gene, a standard curve was established spanning each order of magnitude from 10^1^ to 10^7^ copies of the gene. Target genes were previously obtained by PCR from genomic DNA of *Methanolinea mesofila* (DSMZ 23604) for the *mcrA* gene, and *Methylosinus sporium* (DSMZ 17706) for the gene *pmoA*, both obtained from the DSMZ (German Collection of Microorganisms and Cell Cultures). The qPCR was performed in triplicate for each sample on a StepOne Plus cycler (Thermo Fisher Scientific, Walthman, MA, USA), with a final volume of 10 μL, containing 5 μL of SYBR Green ROX qPCR (Thermo Fisher Scientific, MA, USA), 1 μL of each primer (5 pmols), 1 μL of soil DNA (adjusted to 10 ng/ μL), 0.8 μL of bovine albumin (20 mg / mL) (Sigma-Aldrich, San Luis, MO, USA), and 1.2 μL of ultrapure water (Milli-Q, autoclaved).

In order to minimize bias in the analysis between each qPCR plate run, gene abundance was quantified with the software LinRegPCR (Ramakers et al., 2003). Raw amplification data for each sample were used to calculate individual reaction efficiencies, and detection limits were established for each group of technical replicates. The data generated in arbitrary fluorescence units were converted to the number of copies of the genes using linear interpolation between the known quantities in the standard curve (5 best points out of 7) and the observed fluorescence measurements, using the curves of each plate as a reference for the respective samples.

### Sequencing of 16S rRNA gene fragments

The composition of the microbial community was determined with high throughput sequencing (MiSeq Illumina platform with a 600c kit) of the V4 region of the 16S rRNA gene at the Functional Genomics Center of Luiz de Queiroz College of Agriculture (Caporaso et al., 2011). The V4 region was amplified with the primers 515F (Parada et al., 2016) and 806R (Apprill et al., 2015). This sequencing strategy was selected to match the highly diverse soil environment and for the size of the paired-end reads (average 300 bp). Gene library preparation followed the conditions of 95°C for 3 minutes, followed by 25 cycles at 95°C for 30 seconds, 50°C for 30 seconds, 72°C for 30 seconds, and a final extension step at 72°C for 5 minutes. The DNA concentrations in the samples were adjusted to 10 ng/uL using a Nanodrop 2000c spectrophotometer and the PCR reactions with 2.5 μL of 10x buffer, 1 μL of 50 mM MgCl, 1 µL of 10mM dNTPs, 0.5 μL of 10μM Forward and Reverse Primers, 0.5 μL of 5 U/ μL Taq Platinum - PCR and water for PCR - 14 μL, in a total volume of 25 μL per reaction. Subsequent DNA purification of the amplicon was performed using AMPure XP beads (Beckman Coulter, Brea, CA, USA) and verified on an agarose gel. In a similarly structured, second PCR the adapters were added, followed by another purification with AMPure XP beads and gel electrophoretic confirmation in agarose. The amplicon pool was normalized using quantification by qPCR with the KAPA Illumina quantification kit (Roche, Basel, Switzerland). The computational processing of these data was performed using QIIME2 2017.11 (Bolyen et al., 2019), with data quality control using the DADA2 tool (Callahan et al., 2017), without clustering into OTUs, and taxonomic identification of the sequences was performed using q2-feature-classifier (Bokulich et al., 2018) and the SILVA v.128 99% database (Quast et al., 2013).

### Phylogenetic analyses

Some amplification sequence variants (ASVs) grouped closest with the family Beijerinckiaceae. Since this family represents both methanotrophic and non-methanotrophic genera, we increased our phylogenetic resolution by analyzing phylogenetic trees containing only Beijerinckiaceae sequences that were created with two 16S rRNA primer pairs. One set of primers targeted the region V4, between 515F (Parada et al. 2006) and 806R (Apprill et al. 2015), and a second set of primers targeted the V3/V4 region, between 341F and 805R (Herlemann et al., 2011). For this last pair, the amplification protocol was identical to that described above, except the annealing temperature for the second primer pair was 55°C. The sequences were aligned, and trees were calculated using the software CLC Genomics Workbench 20.0 (QIAGEN, Aarhus, Denmark) at default parameters, and with a maximum likelihood model (PHYML function) with UPGMA (Unweighted Pair Group Method with Arithmetic mean) assuming common replacement frequencies to the bases (Kimura, 1980). The robustness of the final trees was tested with 1000 bootstrap replications. Reference sequences for members of the Beijerinckiaceae family were obtained from the cured database RDP version 11 (Cole et al., 2014) based on the criteria of high-quality reads with a length greater than 1200 bp and representing type strains. The only sequence available for the 16S rRNA gene of the methanotrophic bacterium USCα (Pratscher et al., 2018) was also added.

### Statistical analysis

All comparative analyses between groups were performed with ANOVA followed by a Tukey Honestly Significant Difference (HSD) test and p-values calculated for a two-tailed distribution of the data using the package *agricolae* version 1.2-8 (R Core Team, 2013).

Significant explanatory variables of the methane fluxes were chosen by linear regression and model selection (backward) and by minimizing the Akaike Information Criterion (AIC). The statistical significance was assessed by 1000 permutations of the reduced model. The resulting significant explanatory variables were used to access their contribution to explain the CH_4_ fluxes, using the function *varpart* (Peres-Neto et al., 2016) in the vegan package (Oksanen et al., 2015). Statistical analyses were performed in R Studio software (R Core Team, 2013).

The DEICODE tool (Martino et al., 2019) in the QIIME2 2019.10 was used to process the sequencing data. This tool can identify significant changes in the community based on relative abundance data. Next, the software QURRO (Fedarko et al., 2019) was used to assess shifts in the methane cycling community based on transformed abundance data (natural logarithm) and using a minimum of 10 occurrences per taxon.

## Results

This study tested the effect of acidity correction by liming and the presence of a grass cover by *Urochloa brizantha* cv. Marandu on soil methane fluxes with soils from pasture and forest of different Amazon regions. Two greenhouse experiments were set up. The first experiment was performed with soils from a western Amazon region (Ariquemes, RO) and the second with soils from an eastern region (Tapajós, PA). In both experiments liming resulted in a final pH of ∼6.0 (CaCl_2_, equivalent to pH 6.5 in H_2_O), and an increase in calcium availability, as well a decrease in aluminum saturation (Table S2). In the Ariquemes experiment, methane was sink in both bare soils of forest and pasture at their respective natural pH values, with greater uptake in forest soils (Figure 1). In the Tapajós experiment, we observed methane emissions from bare soils from the pasture at natural pH, and methane uptake in bare soils from forest at natural pH (Figure S1). When forest soils from Ariquemes had grass cover they exhibited the highest methane consumption (Figure 1-b; p = 0.059), at values close to the naturally acidic forest soil, but significantly lower than both limed soils with or without grass cover (Figure 1-b). The Tapajós soils showed a similar trend compared to Ariquemes soils (Figure S1). Methane uptake in pasture soils increase by 35% on average when they have grass coverage (Figure 1-a; p = 0.001). However, liming of pasture soils reduced their methane uptake (Figure 1-a; p = 0.001) and turned forest soils from a methane sink into a methane source (Figure S1-b; p = 0.052).

**Figure 1:**
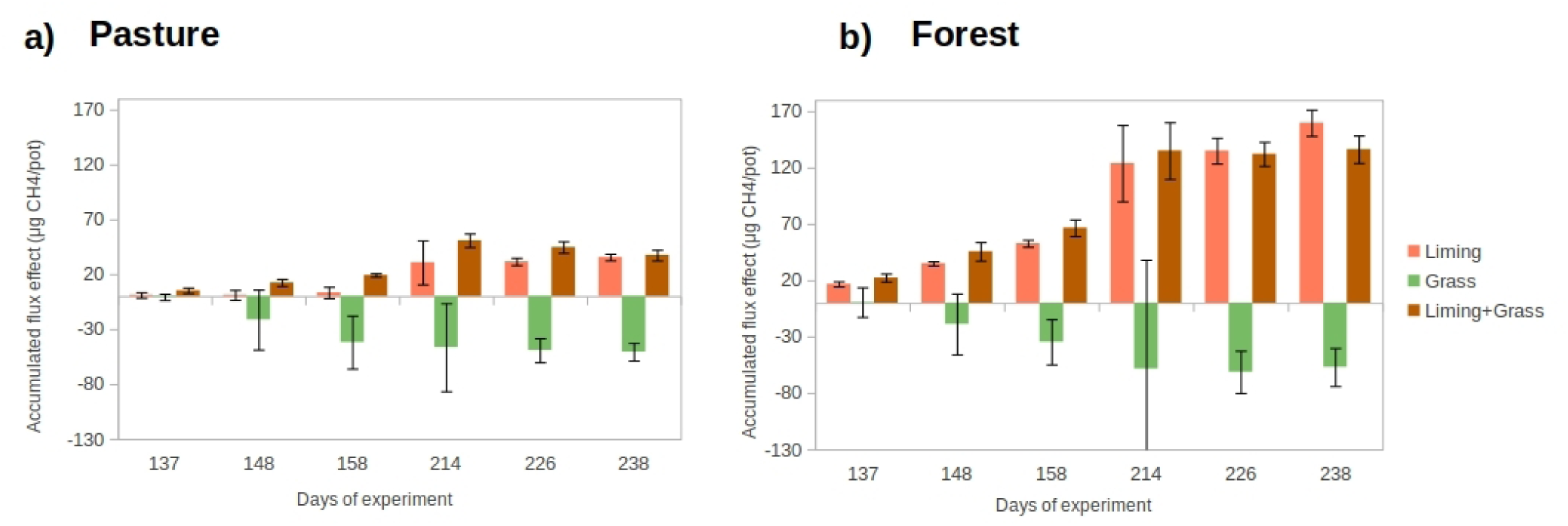
Differences in the accumulated methane flux effect compared to untreated control (bare soils at natural pH) in a) pasture and b) forest soils from the Ariquemes experiment in western Amazonia, with and without acidity correction and with and without grass coverage. Bars show standard deviation.

The validity of our observation that, compared to exposed bare soils, grass coverage improves soil methane uptake was tested in the field. *In situ* measurements of CH_4_ fluxes were taken on two pastures in Belterra/PA, Tapajós region, during the end of wet season, at points with and without grass coverage. No significant differences were observed, but the trend is similar to that observed in the greenhouse experiments (Figure S2; p = 0.112).

Molecular analyses were performed only with soils from the Ariquemes greenhouse experiment (Figure 2) and from the field study in the Tapajós region (Figure S3). The methane cycling microbiota was evaluated through quantification of their functional marker genes *pmo*A and *mcr*A, indicating methane consumers and producers, respectively (Figure 2). The rhizospheric community was evaluated only at the end of the experiment (T3=250 days). During most of the experimental timeline, we did not observe differences in the abundance of methanotrophs between pasture and forest soils (Figure 2). Regarding methane producers, we observed very low abundance in forest soils compared to pasture soils during the experimental duration. The acidity correction shows a tendency to reduce methanotroph levels in forest soils after 250 days in the grass rhizosphere (p = 0.339) and in the bare soil (p = 0.162) (Figure 2A). Pasture soils had between 100 and 1000-fold more methanogenic archaea than forest soils throughout the experiment, which did not change with acidity correction (Figure 2B). The abundance of methanogenic archaea in the grass rhizosphere in pasture soils was reduced on average by 13 times compared to the bare soil (Figure 2B; p = 0.025). No significant changes of methanotrophs were recorded in the rhizosphere (Figure 2A; p = 0.263). This reduction in methanogenic archaea in the grass rhizosphere was not observed in the field study (Figure S3; p = 0.186).

**Figure 2:**
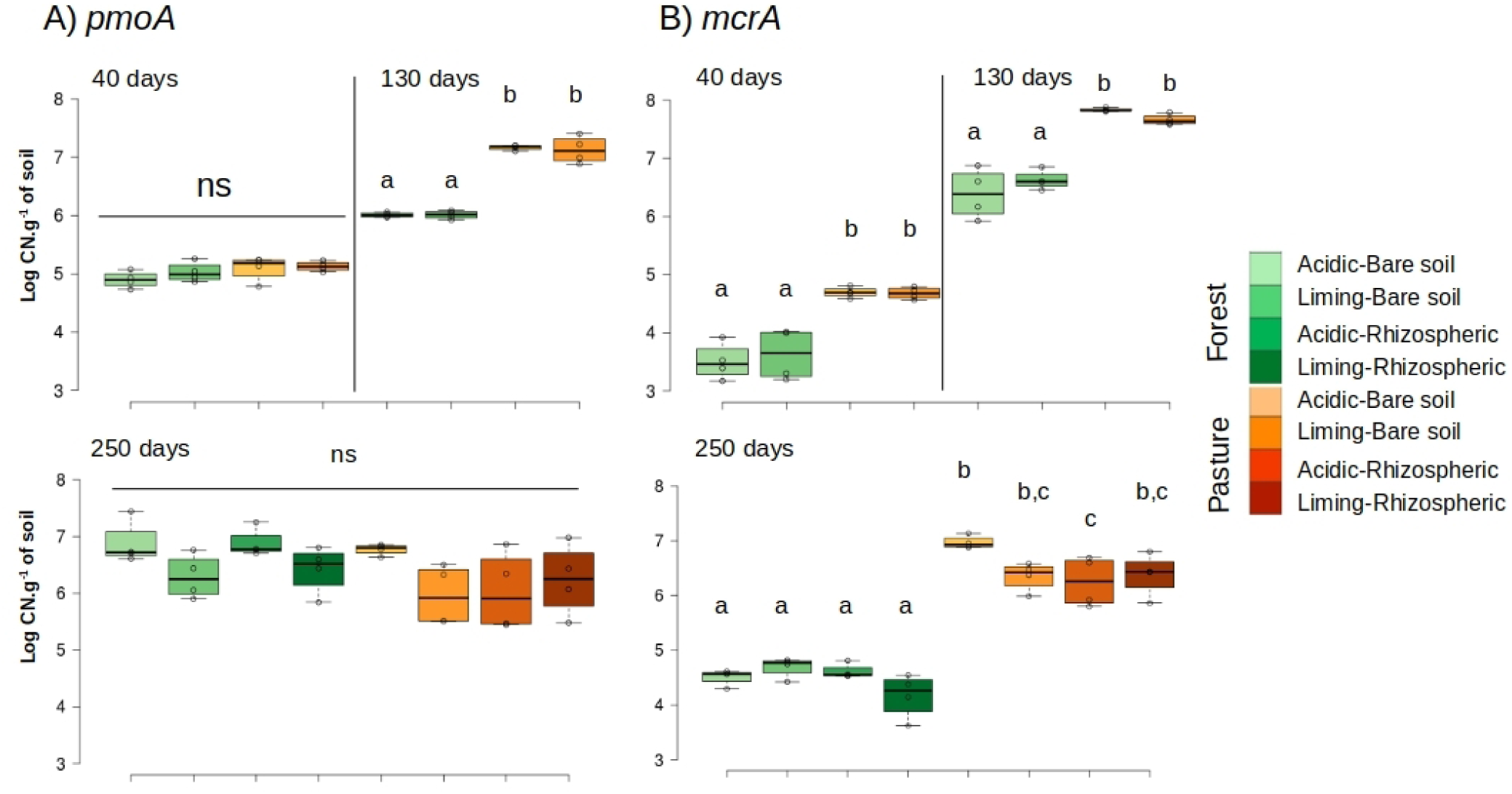
Gene quantification of the genes (A) *pmoA* and (B) *mcrA* at different times after the beginning of the Ariquemes experiment. T1 = 40 days, T2 = 130 days and T3 = 250 days. Letters above each box plot indicate significant changes (p <0.05). CN = copy number. Ns = Not significant.

To investigate the effects of acidity correction and grass rhizosphere on specific groups of microorganisms, high throughput DNA sequencing of the 16S rRNA gene was performed. The results show fair sequencing depth, with rarefaction curves leveling off well below the minimal sequencing depth in soils from the Ariquemes experiment (Table S3, Figure S4-a) and in soils from the field studies (Table S4, Figure S4-b).

Considering only the community associated with methane cycling in the soil (identified with a minimum of 90% confidence), we filtered the groups recognized as methanotrophs (Knief, 2015) and methanogens (Evans et al., 2019) and observed results similar to those obtained in the quantification of gene copies in total DNA. The number of methanogens in forest soils was smaller, while the abundance of methanotrophs was similar between forest and pasture soils (Figure 3). The increase of methanogens in forest soils with acidity correction was not significant. A significant drop in methanotroph abundance was observed only for the combination of acidity correction and grass cover treatments (Figure 3, p=0.024). Methanotrophs in pasture soils did not change with liming or with the presence of grass cover, as previously observed in the quantification of *pmoA* gene copies (Figure 2). However, the number of methanogens was significantly reduced in the grass rhizosphere, with (p=0.017) or without (p=0.007) acidity correction (Figure 3). This last result was similar to that observed in the field, which, although not significant, points to a tendency to reduce methanogenic archaea of different groups in the grass rhizosphere (Figure S5).

**Figure 3:**
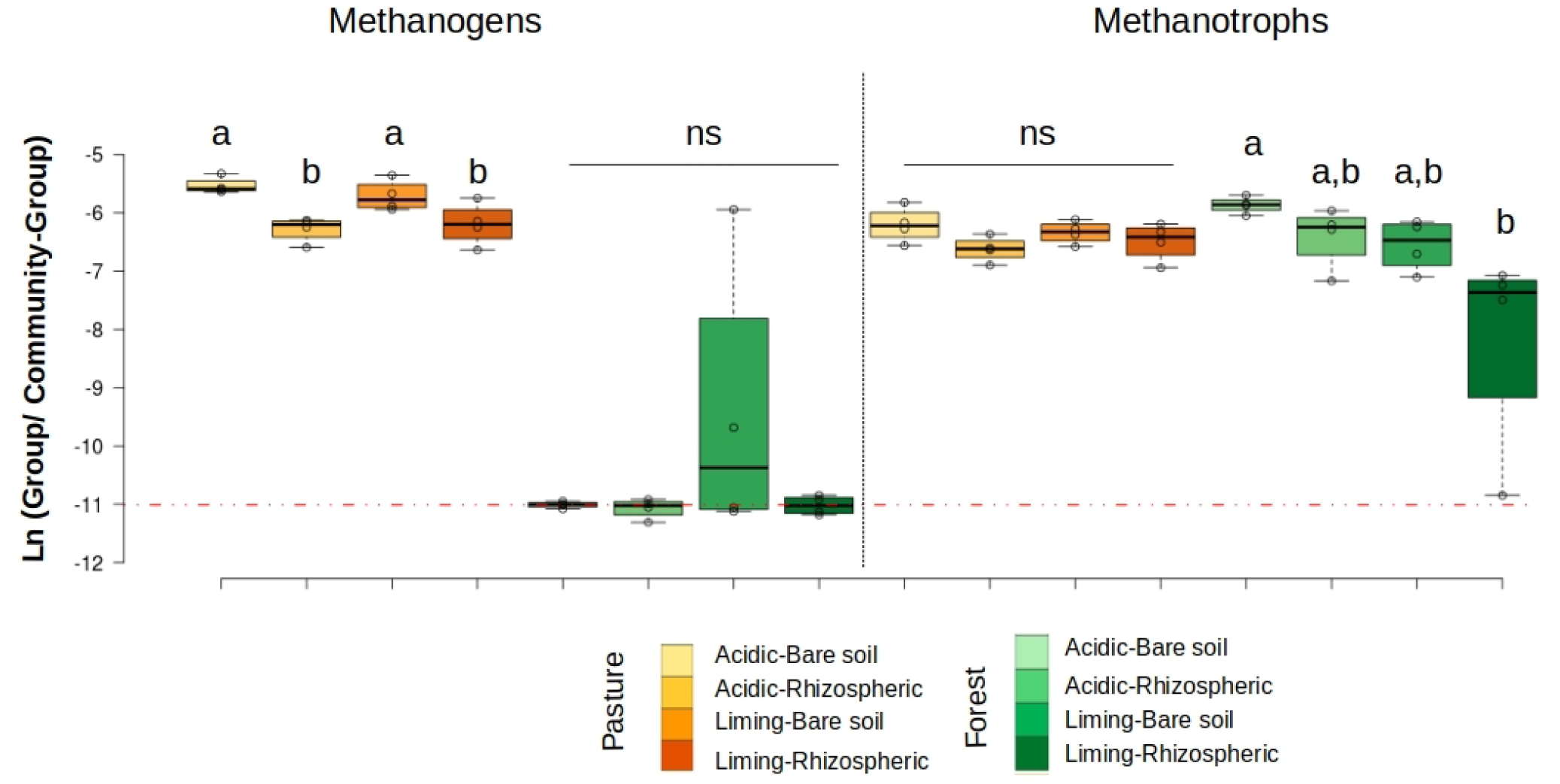
Changes in the logarithmic ratio between methanogenic and methanotrophic groups in relation to the whole community (16S rRNA) in the Ariquemes experiment. The dotted line indicates a calculated ratio of the minimum of 10 counts for each group. The more negative the number of the natural log, the lower the abundance in relation to the total community. Letters indicate significant changes within each treatment for the same land use and group of microorganisms (Tukey HSD; *p*< 0.05). Ns = not significant.

Finally, to understand which groups are associated with methane cycling in these soils, a detailed analysis was performed of all groups that presented sequences of the genera known to act in methane cycling (Knief, 2015; Angel et al., 2012) (Figures 4 and 5). In pasture soils, the abundance of members of all methanogenic genera was lower when soil was grass covered. Forest soils showed a low abundance of methanogenic archaea belonging to *Methanosarcina spp*. The Beijerinckiaceae family is abundant in these soils but it was not possible to identify the sequences at the genus level with the database SILVA v.128 (Quast et al., 2012). A new phylogenetic identification was performed comparing all the sequences annotated as Beijerinckiaceae in the RDP database, with the five amplification sequence variants (ASVs) from sequencing with primers 341F/805R and 8 ASVs from primers 515F/806R. In this analysis, only sequences of the 16S rRNA gene of this family were used as reference, and the results indicate that they are closely related to the methanotrophic clade, with more than 90% confidence (Figures S6 and S7). Those Beijerinckiaceae are reduced in their relative abundance in forest soils with acidity correction (Figure 5), without changes in pasture. No significant changes were observed in relation to other methanotrophs.

**Figure 4:**
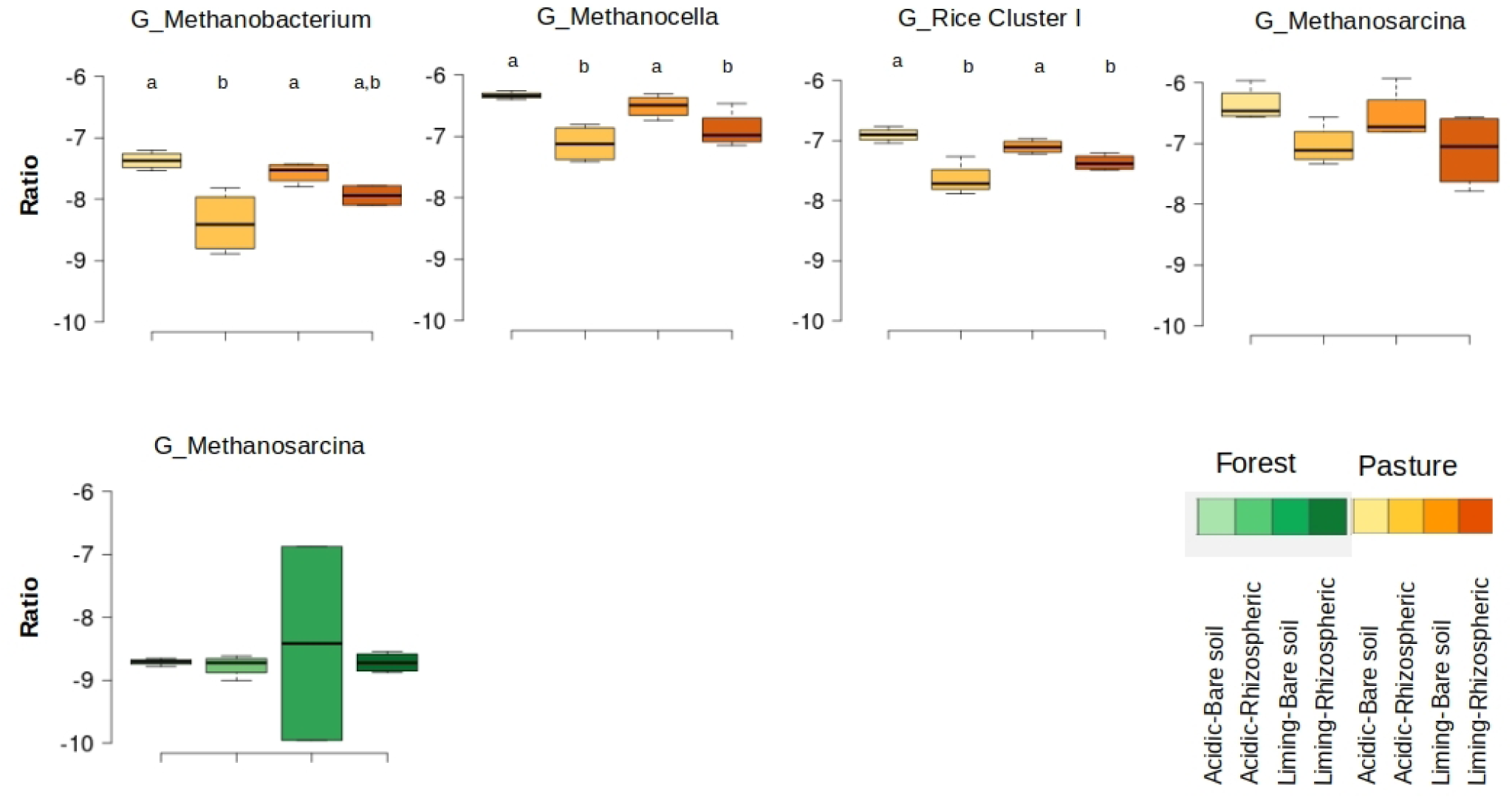
Changes in the Log_e_ ratio between methanogenic microorganisms by genus (G) in relation to the total community in the Ariquemes experiment. All the identified genera are shown. The more negative the numbers, the lower the abundance. Letters indicate significant differences (Tukey HSD; p<0.05)

**Figure 5:**
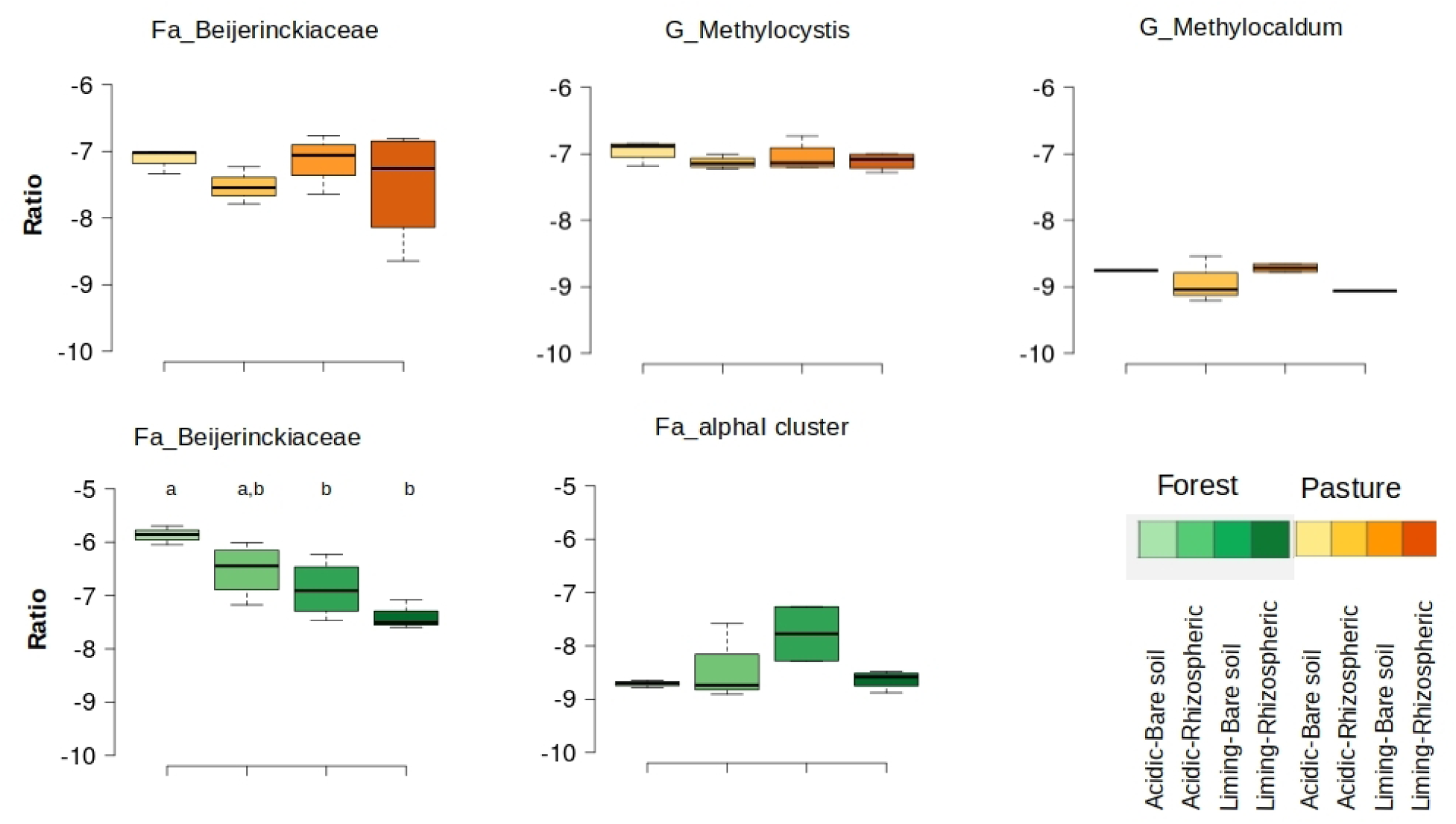
Changes in the Log_e_ ratio between methanotrophic microorganisms by genus (G) or family (Fa) in relation to the total community in the Ariquemes experiment. The more negative the numbers, the lower the abundance. Letters indicate significant differences (Tukey HSD; p<0.05). Analysis showed that a large part of the explained variation (67%) was due to the joint affects.

To disentangle the effect of soil properties, microbial communities, and the presence of grass on the CH_4_ fluxes we performed a variation partitioning analysis. This analysis showed that a large part of the explained variation of the methane fluxes (40%) was due to pH and other soil properties in the first days of the experiment (Figure 6b). However, the grass biomass represented most of the explained variance at the later time-points (Figure 6e-f). The effect of methanogen and methanotroph abundance was minor in the beginning of the incubation (6%) but reached 13% of explained variance at the last time point. These results indicate that the CH_4_ fluxes vary through time and are driven by dynamic factors. Here we decided to separate pH from other soil properties when running this analysis as pH has been previously shown to be a major driver in CH_4_ soil fluxes. Our results confirm this as in the beginning of the experiment the pH and other soil physical-chemical properties were the stronger explanatory variables of the CH_4_ uptake capacity (Table 1). However, the contribution of the soil properties decreases through time while the presence of the grass and the microbial communities gain in explanatory power.

**Table 1.**
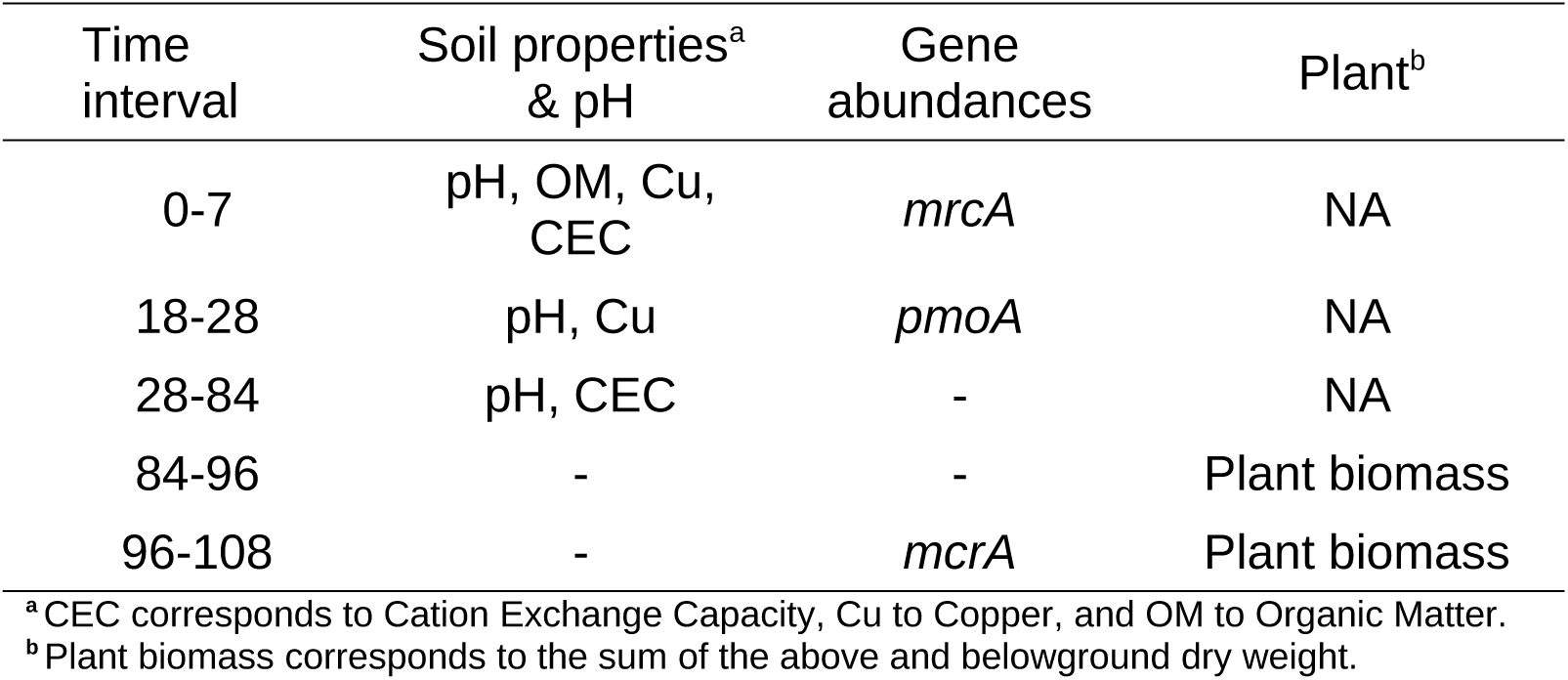
Selected explaining variables for the capacity of methane consumption determined with the variation partitioning analyses at five different time-intervals during the Ariquemes experiment.

| Time interval | Soil properties<sup>a</sup> & pH | Gene abundances | Plant<sup>b</sup> |
| --- | --- | --- | --- |
| 0-7 | pH, OM, Cu, CEC | *mrcA* | NA |
| 18-28 | pH, Cu | *pmoA* | NA |
| 28-84 | pH, CEC | - | NA |
| 84-96 | - | - | Plant biomass |
| 96-108 | - | *mcrA* | Plant biomass |

<sup>a</sup>CEC corresponds to Cation Exchange Capacity, Cu to Copper, and OM to Organic Matter.
<sup>b</sup>Plant biomass corresponds to the sum of the above and belowground dry weight.

**Figure 6:**
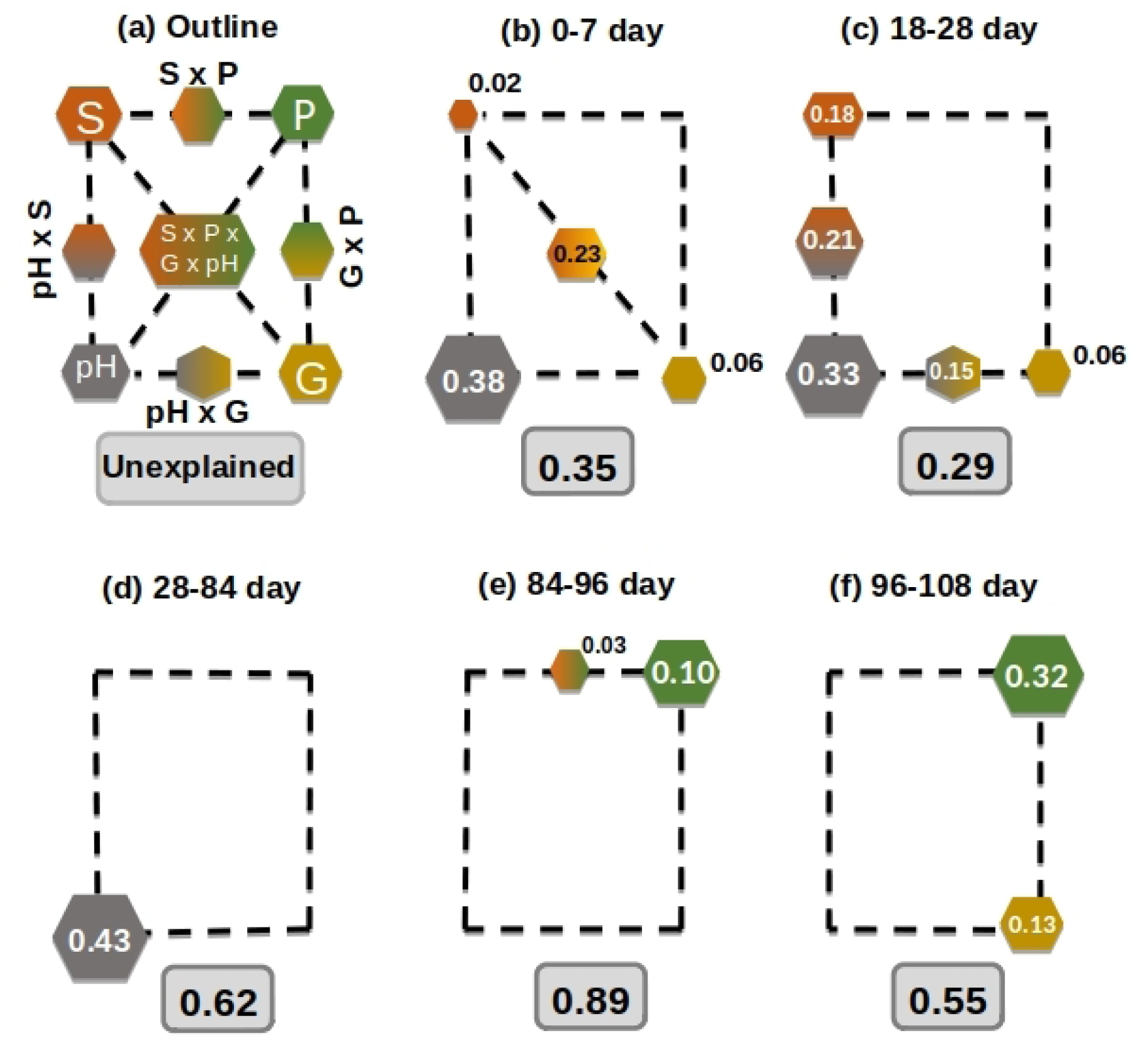
Variation partitioning analysis to determine the drivers of CH_4_ fluxes in the Ariquemes soils in time intervals from day 0 to day 7 (b), day 18 to 28 (c), 28-84 (d), 84-96 (e), and 96-1-8 (f). Variance was partitioned into four explanatory variables, soil physical-chemical properties (S), pH, abundance of methanotrophs and methanogens (G), plant biomass (P), and by combinations of these potential predictors as exemplified in the outline (a). Geometric areas are proportional to the respective percentages of explained variation. The corners of the square depict the variation explained by each factor alone, while percentages of variation explained by interactions of two or all factors are indicated on the sides and in the middle of the square, respectively. All numbers represent percentages, graphically represented by the size of the respective hexagons. Only variance fractions ≥ 2% are shown. The variables used for each variation partitioning are indicated in Table 1.

## Discussion

Deforestation of the Amazonian forest often followed by the establishment of pastures. This forest-to-pasture conversion affects soil methane cycling, where forest soils that were previously acting as a methane sink now become sources of methane (Fernandes et al., 2002). This study confirms previous field results (Meyer et al., 2020; Fernandes et al., 2002; Goreau; De Mello, 1988; Steudler et al., 1996), although the values obtained in our experiments cannot be directly compared to those reported, since the results are limited to a 10 cm surface layer of soil. The trend observed is the same recorded by Steudler et al. (1996), in that forests consume 2.74 more methane than pastures. In our experiments, these values were 0.6-fold in soils from Ariquemes in western Amazonia, and 4.28-fold in soils from Tapajós in eastern Amazonia. This discrepancy may be related to differences in soil microbial communities and chemical properties, but it can also be a consequence of conservation of the forest areas from which these soils originated. While in Ariquemes forests were fragmented and small samples from Tapajós originated from a contiguous forest in a conservation area. Forest fragmentation is known to be associated with increased greenhouse gas emissions (Laurance et al., 1998). Furthermore, the pastures sampled in Ariquemes/RO have a history of long-term management, and the pasture soils sampled in Belterra/PA showed signs of degradation. As management affects the carbon stock in the soil (Fearnside & Imbrozio Barbosa, 1998), it can be expected that it will also affect methane cycling in the soil.

The forest-to-pasture conversion alters the physical-chemical properties of the soil, with consequences to microorganisms that produce and consume methane. In field studies in the same region in Ariquemes, a decrease in methanotrophic bacteria and an increase in methanogenic archaea were observed, in addition to changes in the composition of communities, which were attributed at least in part to changes in methane fluxes (Meyer et al., 2017, Meyer et al. 2020). Also, an increase in activity of methanogens in pastures, compared to forest soils, was recorded in soils from the same region (Kroeger et al. 2020). Our results under controlled moisture conditions did not detect significant changes in the methanotrophic community nor changes in the relative abundance of specific methanotrophic groups. However, there was a significant increase in the abundance of methanogenic archaea in pastures compared to forest soils. These results indicate that studies of the microbiota associated with methane cycling should consider the seasonality of rainfall in the field to better understand the system, such as that developed by Fernandes et al. (2002).

Pasture soils in the Amazonian region present a microbial community quite distinct from that observed in forest soils (de Carvalho et al., 2016; Rodrigues et al., 2013; Jesus et al., 2009). This is partly attributed to acidity reduction in the process of establishing pastures. Forest soils in Amazonia have pH (H_2_O) values between 3.5 to 4.5 (Demattê and Demattê, 1993). The pH is currently understood as one of the main drivers of microbial community structure in soils (Fierer & Jackson, 2006), and increasing it by liming is a strategy to improve fertility and reduce soil toxicity to plants (Oliveira et al., 2013). This process becomes necessary in pasture management to counter the tendency for acidification of pasture soils over time with soil pH reaching values close to those observed in forest areas (de Moraes et al., 1996). In addition, degraded pastures, which can amount to more than 50% of pasture areas in Amazonia (Dias-Filho, 2017), also tend to need acidity correction for their restoration.

Little is known about the effects of soil liming on the methane cycling process in tropical soils. It is known that the optimum growth pH of most cultivable methanotrophs and methanogens is neutral (Le Mer & Roger, 2001; Whittenbury et al., 1970), which is why soil pH represents an important explanatory variable for the distribution of methanotrophs. However, methane oxidation is observed in natural environments across a wide pH range (Knief et al., 2003; Kolb, 2009; Nazaries et al., 2013). Our results indicate that the soil acidity correction for pH (H_2_O) values close to 6.5 has different effects in pasture or forest soils, possibly because the forest soil has undergone a more intense pH correction, starting at 3.5- 5.0 and finishing at 6.5, while in pasture soils the change was from 4.5-5.5 to 6.5. In our forest soils from Ariquemes, we determined a decrease in methane uptake in response to liming, and a shift from uptake to emission in forest soils from Tapajos. Yet, no significant differences were noticed in pasture soils. Thus, liming pasture soils may not impact methane emissions, but still help to maintain the pH of these soils at values suitable for grass biomass productivity. For forest soils, we have shown that the reduction in acidity alone is enough to shift the soil from a methane sink to a source. This change in methane fluxes was not noticeable in the abundance of methanotrophic or methanogenic microorganisms by qPCR, despite a reduction in the relative abundance of methanotrophs that follows the acidity correction.

The identification of microorganisms based on short DNA sequences of the 16S rRNA gene, such as those generated in this study, is limited to the evolutionary information available in that fragment so that it is not always feasible to identify the microorganisms at the genus level. Considering that the ability to oxidize methane is variable at the genus level in the Beijerinckiaceae family, identifying sequences at the family level is not enough to infer if they are methanotrophs. This family also includes generalist bacteria capable of using multiple carbon compounds as an energy source, and here Beijerinckiaceae are more abundant in forest soils than in pastures. Thus, identifying whether they are methanotrophs or not is relevant to understand methane cycling in the forest-to-pasture conversion. The results demonstrate that the Beijerinckiaceae sequences observed in forest soils cluster together in phylogenetic trees. This cluster was observed on the two data sets with high support (> 90% in 1000 bootstraps) and also includes the methanotrophic USCα, which indicates that these sequences are potential methanotrophic Beijerinckiaceae.

The differences observed in methane fluxes after liming were not noticed in the abundance of producers and consumers. This discrepancy might be related to a reduction in the activity of forest soil to act as a methane sink after acidity correction, due to the lower availability of Fe and Cu which are necessary as cofactors for the activity of methane-monooxygenase (Semrau et al., 1995). Alternatively, this discrepancy might be due to limitations of the primers, drawn mostly with microbial references from temperate soils, but we applied them to tropical soils. Or, the difference between methane flux and shift in abundance of methane cycling microorganisms can be due to ammonia oxidizers, possibly oxidizing methane at a higher soil pH. We believe that methanotrophs are the group that was affected the most, since acidity correction was followed by a reduction in the consumption of atmospheric methane by the soil (concentrations of ∼1.8 ppm). Also, the duration of incubating soil from Ariquemes for 250 days should be long enough to observe compensatory changes due to DNA replication, that should be detected in the DNA quantification analysis.

Although there is great natural variability in the methane flux data, the final averages led to the conclusion that pasture soils act as a methane source, which is in fact a commonly reported final result. This observed variability means that pastures could seasonally or by location switch from being a methane source to temporarily becoming a methane sink (Fernandes et al., 2002; Steudler et al., 1996). Our initial hypothesis was that the methane consuming capacity of pasture soils would be related to intermediate moisture availability in the micro-environments of soil, since soil moisture is a determining factor for methane fluxes in pastures (Verchot et al., 2000). To eliminate moisture variation as a variable in the experiments we set the soil water contents at 70% of the holding capacity in the greenhouse experiments. The variability of pasture gas fluxes could also be explained by grass coverage, a factor associated with pasture management. The management of pastures can influence soil gas fluxes (Figueiredo et al. 2017), since it influences the carbon stocks in the soil (Fearnside & Imbrozio Barbosa, 1998), however the way ongoing pasture management can affect the microbial community remains an open question. Considering that management is performed with the goal of grass productivity, and greater aerial biomass is associated with greater root biomass, we expect that a larger root surface area in pasture would create a more interactive environment with the soil microbiota, and thus enable higher rhizosphere activity. The role of the rhizosphere on methane cycling in upland soils is still poorly understood, and even different plant species can influence the soil by increasing methane oxidation or production, depending on the type of soil or soil conditions (Praeg et al., 2017). In soils of the Ariquemes experiment, we observed that plant cover will lead to a reduced methane flux in both forest and pasture soils compared to those with acidity correction. The methane flux rates with grass cover were similar to those of the original forest soil and tended to be higher than those of pasture without acidity correction. In soils from Tapajós experiment, the same trend was observed in forest soils, but possible due to the shorter duration of this experiment, there were no significant differences in the pasture.

When disentangling the contribution of different biotic and abiotic factors to CH_4_ soil uptake capacity we found that its drivers change through time, which could explain as previously discussed that soils might change from a source to a sink. While pH and other soil properties explained most of the variance in the beginning of the greenhouse experiment, the abundance of microbial communities related to CH_4_ fluxes and plant biomass explained most of the CH_4_ uptake at the end of the experiment. These results suggests that our treatments (liming and planting grass) are causing a reorganization of the microbial communities and while the soil properties are initially the main variables explaining the CH_4_ fluxes, after a couple of weeks the biotic factors are the main drivers of CH_4_ fluxes in these soils. While a previous study showed that peak emissions of the green-house gas N_2_O can be driven by the microorganisms related to the production and reduction of this green-house gas (Domeignoz-Horta, et al., 2017), our results show how microorganisms related to methane cycling and plant cover play a role to understand the temporal dynamics of CH_4_ uptake in soils. These results highlight the need for better characterizing microbial communities to increase our understanding of the relationship between abundance and diversity of microorganisms and their corresponding processes.

The results presented here demonstrate that soil acidity is an important factor for methane sequestration in tropical soils, as the acidity correction reduces this capacity. In pastures, the effect of the acidity correction is less consequential compared to the presence of grass coverage. This demonstrates that the correction of acidity in pastures, if combined with constant soil coverage with grass, would have little or no impact on methane emissions while improving soil structure and increasing nutrient availability, soil organic matter and grass productivity.

## Conclusion

Our results suggest that the impact of methane emissions from pastures in the Amazonian region can be mitigated through pasture management, specifically by keeping the soil always covered with grass. The rhizosphere of *Urochloa brizantha* cv. Marandu affects soil microbial communities by lowering the abundance of methanogenic archaea up to 10 times compared to the bare soil. The affected methanogens are composed of *Methanobacterium spp*., *Methanocella spp*., *Rice Cluster I, and Methanosarcina spp*. In addition, we demonstrate that the correction of acidity in pasture soils can reduce methane sequestration under atmospheric methane concentrations (high-affinity methanotrophs). Therefore, the level of acidity correction should be considered as a factor for additional emissions of greenhouse gases. In the acidic forest soils, an increase in pH reduced methane sequestration by more than 50%, thereby reversing the flux direction to turn forest soil from a methane sink into a source. Field studies with liming and a focus on the grass rhizosphere under seasonal conditions are urgently needed to provide specific recommendations to policymakers and farmers.

## Acknowledgments

The authors thank the owners and staff of Farm “Fazenda Nova Vida”, for logistical support and permission to work on their property. We also thank the private landowners: Aristeu, Bernardo e Elói for their support and access to their land. We would like to thank the Large-Scale Biosphere-Atmosphere Program (LBA), coordinated by the National Institute for Amazon Research (INPA), for the use and availability of data for logistical support and infrastructure during field activities. Additionally, we are grateful to Prof. Plinio B. de Camargo, Henrique Cipriani (EMBRAPA-RO), Alexandre Pedrinho, and to Wagner Piccinini for assistance with fieldwork.

## Funding

This project was supported by the BIOTA FAPESP and NSF – Dimensions of Biodiversity (2014/50320-4 and DEB 1442183) and by CNPq (311008/2016-0). Additional funding in the form of scholarships were provided by FAPESP (2018/09117-1), CNPq (140953/2017-5), and CAPES (001 and 88881.189492/2018-01).

## Notes

### Competing Interest Statement

The authors have declared no competing interest.

