## Supplementary Material for "Maintaining grass coverage increases methane uptake in Amazonian pasture soils"

Table S1: Description of the primer sets used in the qPCR assays, targeting the marker genes mcrA and pmoA.

| **Gene** | **Primer** | **Sequence (5’-3’)** | **Cycling Parameters** | **Reference** |
| --- | --- | --- | --- | --- |
| *mcrA* | mlas-F | GGYGGTGTMGGDTTCACMCARTA | 95ºC for 10’ –  45 cycles: 95ºC for 30” and 60ºC for 45” and 72ºC for 30” | Angel et al. (2012) |
|  | mcrA-R | CGTTCATBGCGTAGTTVGGRTAGT |  | Steinberg and Regan (2008) |
| *pmoA* | A189F | GGNGACTGGGACTTCTGG | 95ºC for 10’ –  45 cycles: 95ºC for 30” and 58ºC for 30” and 72ºC for 45” | Holmes et al. (1999) |
|  | MB661r | CCGGMGCAACGTCYTTACC |  | Costello and Lidstrom (1999) |

Table S2: Soil chemical attributes for different times and treatments of the experiments (continued)

| Area | Time | Treatment | pH | OM | P | K | Ca | Mg | H+Al | Al |
| --- | --- | --- | --- | --- | --- | --- | --- | --- | --- | --- |
|  |  |  | **CaCl_2_** | **g.dm^-3^** | **mg.dm^-3^** | **mmolc.dm^-3^** | **mmolc.dm^-3^** | **mmolc.dm^-3^** | **mmolc.dm^-3^** | **mmolc.dm^-3^** |
| Ariquemes/RO | Day 0 | Forest – natural pH – bare soil | 4.03 ± 0.06 | 28.3 ± 1.1 | 9.0 ± 0.0 | 1.2 ± 0.0 | 5.3 ± 1.1 | 3.7 ± 0.6 | 38.0 ± 4.0 | 6.0 ± 0.0 |
|  |  | Pasture-natural pH-bare soil | 4.63 ± 0.06 | 34.0 ± 2.0 | 7.3 ± 1.1 | 1.6 ± 0.3 | 11.3 ± 3.5 | 4.3 ± 1.1 | 29.0 ± 1.7 | <2 |
|  | Day 250 | Forest-natural pH-bare soil | 4.20 ± 0.18 | 27.7 ± 3.0 | 5.5 ± 1.0 | 1.2 ± 0.9 | 6.2 ± 1.7 | 1.7 ± 0.5 | 61.5 ± 8.5 | 6.5 ± 0.6 |
|  |  | Forest-natural pH-grass | 4.20 ± 0.12 | 26.0 ± 2.0 | 5.0 ± 0.8 | 0.9 ± 0.6 | 6.5 ± 2.1 | 2.2 ± 0.5 | 50.0 ± 8.5 | 6.0 ± 0.8 |
|  |  | Forest-liming-bare soil | 6.15 ± 0.17 | 24.0 ± 1.1 | 8.5 ± 2.6 | 0.7 ± 0.2 | 42.5 ± 3.7 | 2.7 ± 0.5 | 14.0 ± 1.1 | <2 |
|  |  | Forest-liming-grass | 6.05 ± 0.17 | 25.0 ± 1.6 | 8.5 ± 1.9 | 0.6 ± 0.1 | 38.2 ± 3.7 | 2.0 ± 0.0 | 14.7 ± 1.3 | <2 |
|  |  | Pasture-natural pH-bare soil | 4.67 ± 0.15 | 31.7 ± 3.3 | 6.2 ± 2.6 | 1.4 ± 0.1 | 15.2 ± 1.5 | 5.7 ± 0.5 | 37.0 ± 2.0 | <2 |
|  |  | Pasture-natural pH-grass | 4.75 ± 0.17 | 26.5 ± 1.0 | 5.5 ± 0.6 | 1.2 ± 0.0 | 15.5 ± 1.9 | 6.0 ± 0.0 | 34.0 ± 0.0 | <2 |
|  |  | Pastage-liming-bare soil | 5.92 ± 0.09 | 27.7 ± 3.0 | 8.2 ± 3.2 | 1.3 ± 0.1 | 36.7 ± 6.5 | 5.2 ± 0.5 | 15.5 ± 0.6 | <2 |
|  |  | Pasture-liming-grass | 5.97 ± 0.15 | 30.7 ± 2.5 | 8.2 ± 2.4 | 0.9 ± 0.1 | 42.0 ± 7.5 | 4.2 ± 1.0 | 15.5 ± 2.1 | <2 |
| Belterra/PA | Day 0 | Forest-natural pH-bare soil | 3.70 ± 0.30 | 39.6 ± 10.3 | 6.2 ± 1.3 | 1.1 ± 0.2 | 1.6 ± 0.5 | 2.4 ± 0.5 | 143 ± 44.6 | 16.8 ± 1.6 |
|  |  | Pasture-natural pH-bare soil | 4.50 ± 0.10 | 31.8 ± 3.0 | 3.6 ± 1.1 | 1.1 ± 0.2 | 12.2 ± 1.9 | 7.2 ± 1.5 | 44 ± 2.7 | 2.2 ± 0.8 |

O.M. = Organic Matter; H+Al = Potential acidity; A = Natural pH; B = Bare soil; L = Liming; G = Grass.

Table S2: Soil chemical attributes for different times and treatments of the experiments (conclusion)

| Area | Time | Treatment | BS | CEC | V | m | S | Cu | Fe | Mn | B |
| --- | --- | --- | --- | --- | --- | --- | --- | --- | --- | --- | --- |
|  |  |  | **mmolc.dm^-3^** | **mmolc.dm^-3^** | **%** | **%** | **mg.dm^-3^** | **mg.dm^-3^** | **mg.dm^-3^** | **mg.dm^-3^** | **mg.dm^-3^** |
| Ariquemes/RO | Day 0 | Forest – natural pH – bare soil | 10.0 ± 1.0 | 48.0 ± 3.3 | 21.0 ± 3.5 | 27.6 ± 15.4 | 11.0 ± 2.8 | 0.5 ± 0.1 | 83.3 ± 7.8 | 6.5 ± 2.5 | <0.15 |
|  |  | Pasture-natural pH-bare soil | 17.3 ± 4.7 | 46.3 ± 5.0 | 37.0 ± 6.6 | 6.0 ± 7.2 | <6 | 1.3 ± 0.1 | 84.3 ± 7.5 | 78.2 ± 8.7 | <0.15 |
|  | Day 250 | Forest-natural pH-bare soil | 9.5 ± 2.9 | 71.0 ± 7.3 | 13.2 ± 4.6 | 42.5 ± 9.9 | 7.7 ± 2.4 | 0.3 ± 0.0 | 133.8 ± 42.4 | 4.9 ± 0.9 | <0.15 |
|  |  | Forest-natural pH-grass | 10.0 ± 2.4 | 60.0 ± 6.8 | 16.5 ± 5.7 | 39.0 ± 8.1 | <6 | 0.3 ± 0.1 | 105.0 ± 22.0 | 3.2 ± 0.5 | 0.2 ± 0.1 |
|  |  | Forest-liming-bare soil | 46.2 ± 4.1 | 60.2 ± 3.1 | 76.5 ± 3.0 | ns | <6 | 0.2 ± 0.0 | 20.2 ± 5.7 | 1.5 ± 0.6 | 0.3 ± 0.2 |
|  |  | Forest-liming-grass | 41.2 ± 3.7 | 56.0 ± 2.6 | 73.5 ± 3.3 | ns | 6.5 ± 1.3 | 0.2 ± 0.1 | 27.6 ± 6.7 | 1.8 ± 0.4 | 0.2 ± 0.1 |
|  |  | Pasture-natural pH-bare soil | 22.7 ± 2.0 | 59.7 ± 1.5 | 37.7 ± 3.1 | 8.5 ± 0.6 | <6 | 1.2 ± 0.2 | 108.0 ± 11.6 | 34.5 ± 10.4 | <0.15 |
|  |  | Pasture-natural pH-grass | 22.5 ± 1.9 | 56.5 ± 1.9 | 40.0 ± 2.4 | 6.5 ± 2.9 | 7.2 ± 3.4 | 1.1 ± 0.1 | 103.4 ± 9.8 | 34.5 ± 4.9 | 0.2 ± 0.1 |
|  |  | Pastage-liming-bare soil | 43.2 ± 6.9 | 58.7 ± 6.7 | 73.5 ± 3.7 | ns | <6 | 0.8 ± 0.2 | 33.6 ± 6.3 | 14.3 ± 4.2 | <0.15 |
|  |  | Pasture-liming-grass | 47.2 ± 8.4 | 62.7 ± 7.8 | 74.7 ± 4.9 | ns | <6 | 0.8 ± 0.1 | 30.8 ± 7.9 | 17.4 ± 3.5 | <0.15 |
| Belterra/PA | Day 0 | Forest-natural pH-bare soil | 5 ± 1.0 | 148 ± 44.9 | 3.4 ± 1.1 | 76.6 ± 4.9 | 11 ± 2.5 | 0.3 ± 0.1 | 161.6 ± 127.3 | 9.2 ± 7.9 | 0.5 ± 0.1 |
|  |  | Pasture-natural pH-bare soil | 20.6 ± 3.7 | 64.6 ± 4.6 | 31.8 ± 3.9 | 10.2 ± 4.8 | 11.4 ± 1.5 | 0.4 ± 0.1 | 151.8 ± 52.1 | 4.8 ± 1.2 | 0.5 ± 0.1 |

BS = Bases Sum; CEC = Cation Exchange Capacity; V = Bases saturation; m = Aluminum saturation; A = Natural pH; B = Bare soil; L = Liming; G = Grass

Table S3: Sequencing depth from samples of experiment 1 (Ariquemes/RO - 2017), from amplicons after filtering, denoising, fusion of forward and reverse sequences, and removal of chimeras using the software DADA2. Lower value is 50,993 ASVs.

| **Sample code** | **Treatment** | **ASVs after DADA2** |
| --- | --- | --- |
| FP 0-0-1 | FP-natural pH-bare soil | 64909 |
| FP 0-0-2 | FP-natural pH-bare soil | 59810 |
| FP 0-0-3 | FP-natural pH-bare soil | 56995 |
| FP 0-0-4 | FP-natural pH-bare soil | 60649 |
| FP 0-1-1 | FP-natural pH-grass | 59535 |
| FP 0-1-2 | FP-natural pH-grass | 63362 |
| FP 0-1-3 | FP-natural pH-grass | 55209 |
| FP 0-1-4 | FP-natural pH-grass | 81738 |
| FP 1-0-1 | FP-liming-bare soil | 73568 |
| FP 1-0-2 | FP-liming-bare soil | 63641 |
| FP 1-0-3 | FP-liming-bare soil | 64302 |
| FP 1-0-4 | FP-liming-bare soil | 67549 |
| FP 1-1-1 | FP-liming-grass | 71862 |
| FP 1-1-2 | FP-liming-grass | 55383 |
| FP 1-1-3 | FP-liming-grass | 51441 |
| FP 1-1-4 | FP-liming-grass | 68334 |
| P72 0-0-1 | Pasture-natural pH-bare soil | 83421 |
| P72 0-0-2 | Pasture-natural pH-bare soil | 70108 |
| P72 0-0-3 | Pasture-natural pH-bare soil | 78405 |
| P72 0-0-4 | Pasture-natural pH-bare soil | 63618 |
| P72 0-1-1 | Pasture-natural pH-grass | 70628 |
| P72 0-1-2 | Pasture-natural pH-grass | 50993 |
| P72 0-1-3 | Pasture-natural pH-grass | 82473 |
| P72 0-1-4 | Pasture-natural pH-grass | 74254 |
| P72 1-0-1 | Pasture-liming-bare soil | 95860 |
| P72 1-0-2 | Pasture-liming-bare soil | 88514 |
| P72 1-0-3 | Pasture-liming-bare soil | 69433 |
| P72 1-0-4 | Pasture-liming-bare soil | 58978 |
| P72 1-1-1 | Pasture-liming-grass | 63146 |
| P72 1-1-2 | Pasture-liming-grass | 61570 |
| P72 1-1-3 | Pasture-liming-grass | 69874 |
| P72 1-1-4 | Pasture-liming-grass | 77523 |

P = Pasture, FP = Primary Forest.

Table S4: Sequencing depth from samples of the field study (Belterra/PA - 2019), from amplicons after filtering, denoising, fusion of forward and reverse sequences, and removal of chimeras using the software DADA2. Lower value is 11,761 ASVs. P = Pasture.

| Sample | Treatment | ASVs number after DADA2 |
| --- | --- | --- |
| B-1-A-2 | P1-Adjacent to the Roots | 40675 |
| B-1-B-3 | P1-bare soil | 46030 |
| B-1-R-1 | P1-Rhizosphere | 25993 |
| B-3-A-8 | P1-Adjacent to the Roots | 86869 |
| B-3-B-9 | P1-bare soil | 46584 |
| B-3-R-7 | P1-Rhizosphere | 52615 |
| B-4-A-11 | P1-Adjacent to the Roots | 23657 |
| B-4-B-12 | P1-bare soil | 20374 |
| B-4-R-10 | P1-Rhizosphere | 65066 |
| B-5-A-14 | P1-Adjacent to the Roots | 21783 |
| B-5-B-15 | P1-bare soil | 22187 |
| B-5-R-13 | P1-Rhizosphere | 25661 |
| G-1-A-17 | P2-Adjacent to the Roots | 38360 |
| G-1-B-18 | P2-bare soil | 42145 |
| G-1-R-16 | P2-Rhizosphere | 31777 |
| G-3-A-23 | P2-Adjacent to the Roots | 26987 |
| G-3-B-24 | P2-bare soil | 15246 |
| G-3-R-22 | P2-Rhizosphere | 30297 |
| G-4-A-26 | P2-Adjacent to the Roots | 21832 |
| G-4-B-27 | P2-bare soil | 14162 |
| G-4-R-25 | P2-Rhizosphere | 20555 |
| G-5-A-29 | P2-Adjacent to the Roots | 11761 |
| G-5-B-30 | P2-bare soil | 19760 |
| G-5-R-28 | P2-Rhizosphere | 17842 |

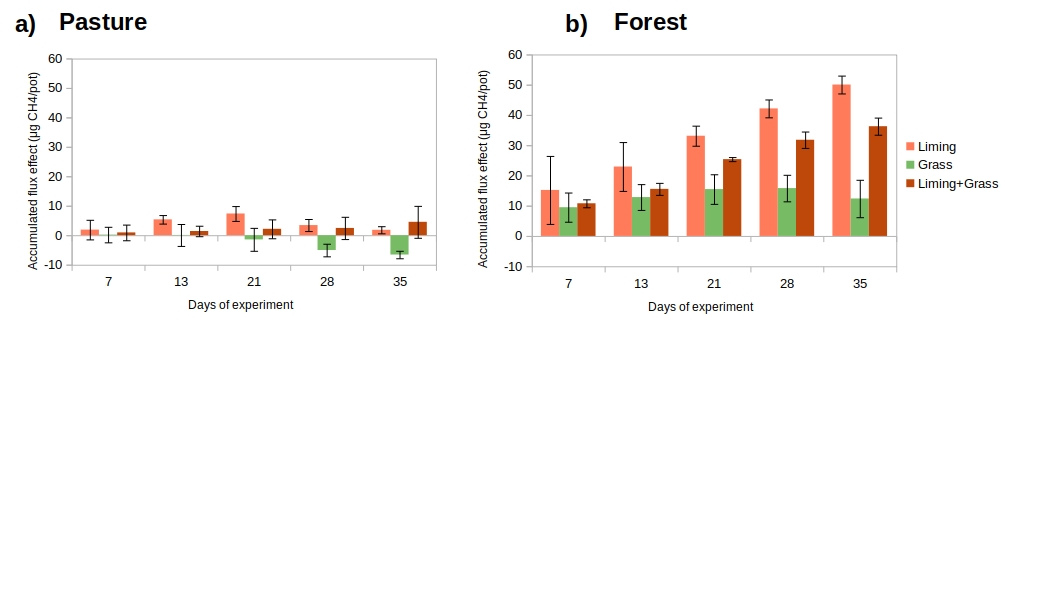
Figure S1: Differences in the cumulative CH_4_ fluxes discounted the control soils (bare natural pH soils) in a) pasture and b) forest soils from the Tapajós experiment in eastern Amazonia, with and without acidity correction and with and without grass coverage. Bars show standard deviation.

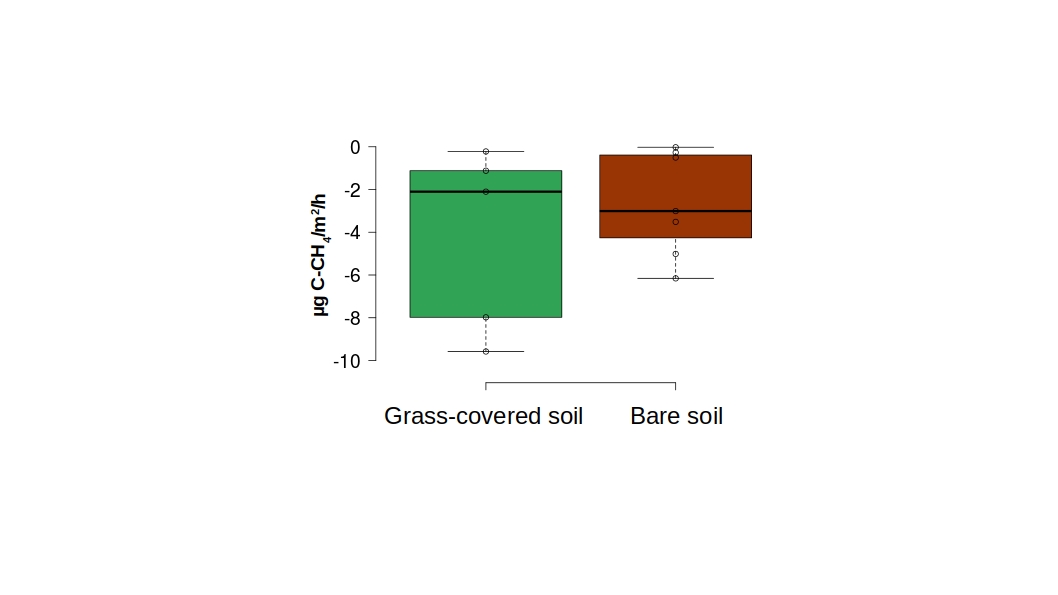

Figure S2: CH_4_ flux in the field in pasture areas from Belterra, PA – Tapajós region, in eastern Amazonia under grass cover and in areas without grass coverage (bare soil). There is no significant difference between the two groups (p=0,11).

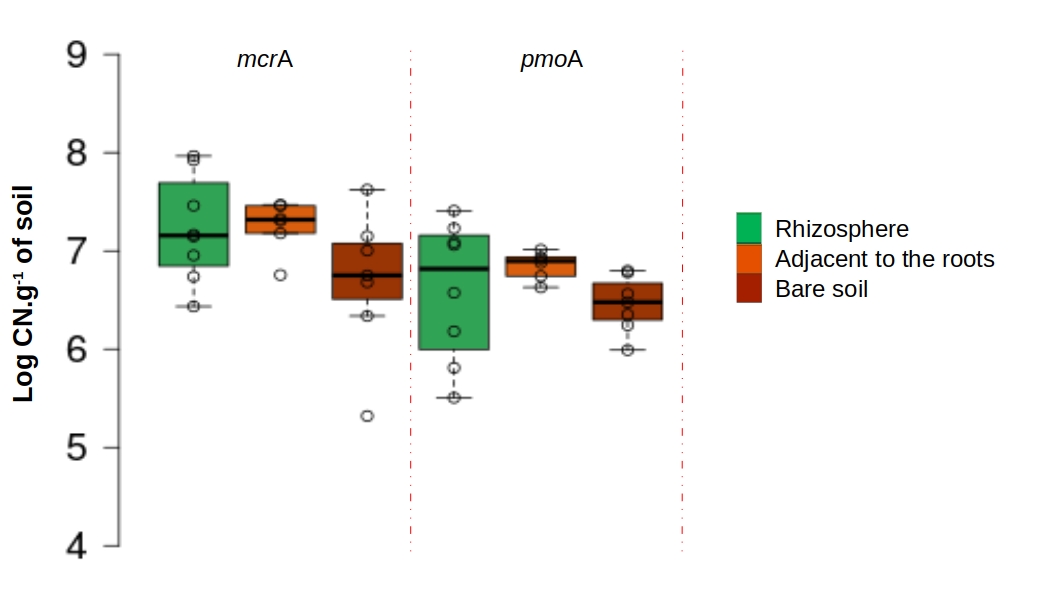

Figure S3: Quantification of gene copies of methanogens (*mcrA*) and methanotrophs (*pmoA*) from field samples of pastures in Belterra, PA – Tapajós region. The grass rhizosphere, the root zone (soils adjacent to the plant, but not attached to the root), and bare pasture areas (bare soil without grass cover) were compared. No significant differences were found.

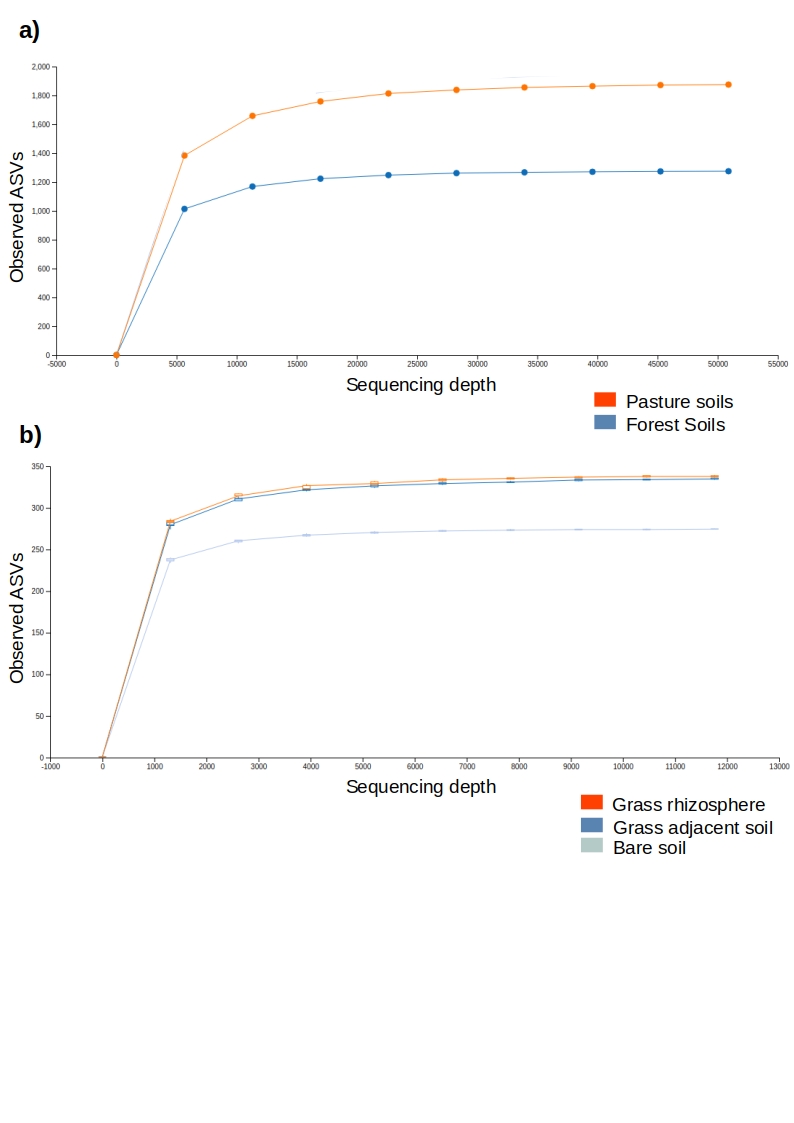

Figure S4: Rarefaction analysis of sequencing data created from soil extracted DNA from a) greenhouse experiments with soils from the Ariquemes experiment and from b) field studies of pastures (Belterra, PA – Tapajós region).

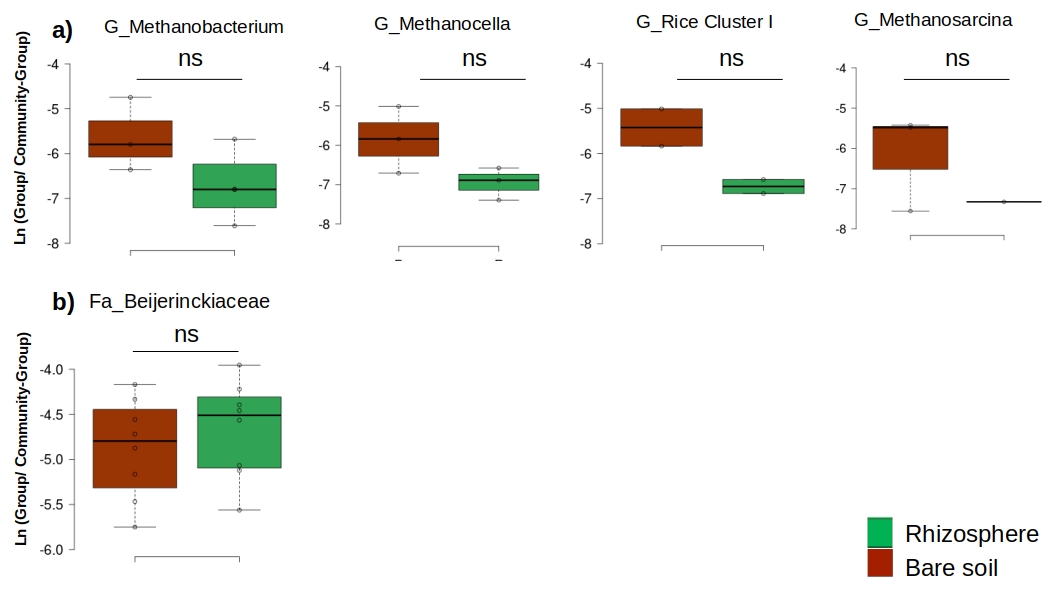

Figure S5: Log_10_ ratio between (a) methanogenic microorganisms and (b) methanotrophic microorganisms by genus (G) or family (Fa) in relation to the total community. Field samples of pasture areas from Belterra, PA – Tapajós region – are shown where either the grass rhizosphere soil or bare soil were used. The more negative the numbers, the lower the abundance. ns = not significant.

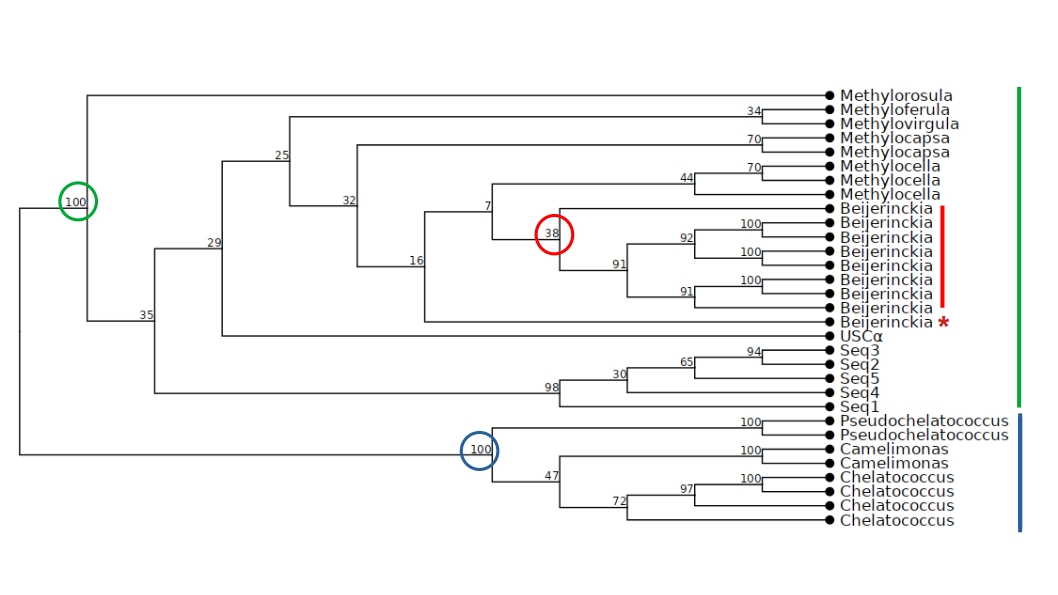
Figure S6: Phylogenetic tree based on 16S rRNA sequences and identified as Beijerinckiaceae (Seq1 to Seq5), along with sequences available in the database for microorganisms of this family and of USCα. Values displayed show percentage of support for groupings after 1000 bootstraps. Non-methanotrophs are indicated by the blue bar, methanotrophs together with our sequences are in green and the genus *Beijerinckia* known for a secondary loss of methane oxidation capacity is in red (Tamas et al. 2014). * indicates ambiguous positions.

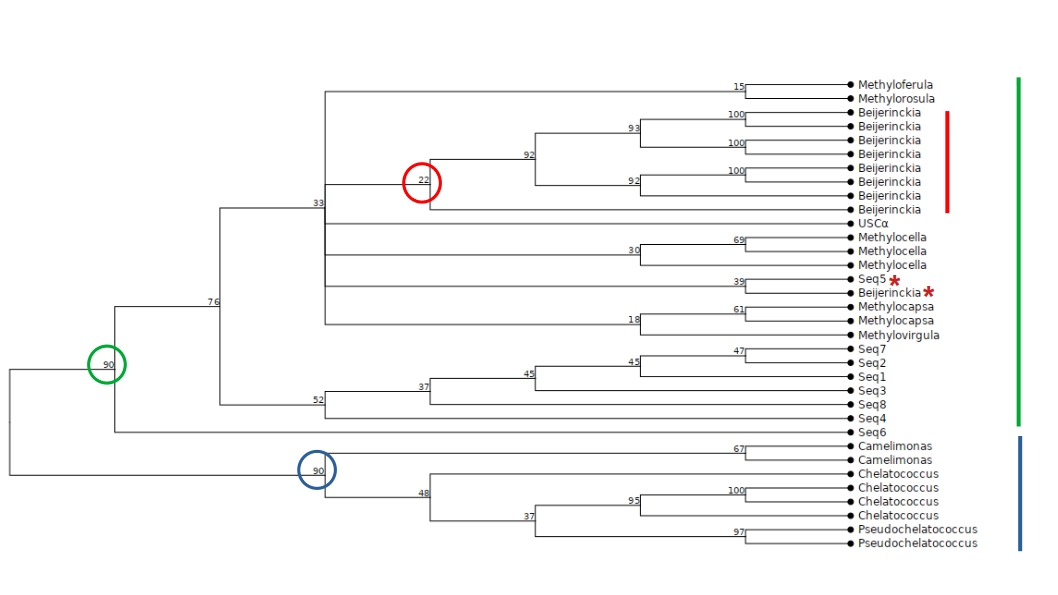
Figure S7: Phylogenetic tree with sequences of the 16S rRNA gene obtained from the primer pair 515F/806R for region V4 and identified as Beijerinckiaceae (Seq1 to Seq8), along with the high-quality type sequences available for microorganisms of this family and of USCα. Values displayed show percentage of support for groupings after 1000 bootstraps. Non-methanotrophs are indicated by the blue bar, methanotrophs together with our sequences are in green and the genus *Beijerinckia* known for a secondary loss of methane oxidation capacity is in red (Tamas et al. 2014). * indicates ambiguous positions.
